# Two-antibody pan-ebolavirus cocktail confers broad therapeutic protection in ferrets and nonhuman primates

**DOI:** 10.1101/395533

**Authors:** Zachary A. Bornholdt, Andrew S. Herbert, Chad E. Mire, Shihua He, Robert W. Cross, Anna Wec, Dafna M. Abelson, Joan B. Geisbert, Rebekah M. James, Md Niaz Rahim, Wenjun Zhu, Viktoriya Borisevich, Logan Banadyga, Bronwyn M. Gunn, Krystle N. Agans, Eileen Goodwin, Kevin Tierney, William S. Shestowsky, Ognian Bohorov, Natasha Bohorova, Jesus Velasco, Eric Ailor, Do Kim, Michael H. Pauly, Kevin J. Whaley, Galit Alter, Laura M. Walker, Kartik Chandran, Larry Zeitlin, Xiangguo Qiu, Thomas W. Geisbert, John M. Dye

**Affiliations:** Mapp Biopharmaceutical, Inc. San Diego, CA, USA; United States Army Medical Research Institute of Infectious Diseases, Fort Detrick, MD, USA; Department of Microbiology and Immunology, Galveston National Laboratory, University of Texas Medical Branch at Galveston, TX, USA; Special Pathogens Program, National Microbiology Laboratory, Public Health Agency of Canada, Winnipeg, MB, Canada; Department of Microbiology and Immunology, Albert Einstein College of Medicine, Bronx, NY, USA; The Ragon Institute of MGH, MIT, and Harvard, Cambridge, MA, USA; Department of Medical Microbiology, University of Manitoba, Winnipeg, MB, Canada; Adimab LLC, Lebanon, NH, USA

## Abstract

All available experimental vaccines and immunotherapeutics^1,2^ against Ebola virus (EBOV), including rVSV-ZEBOV^3^ and ZMapp^TM4^, lack activity against other ebolaviruses associated with human disease outbreaks. This year, two separate outbreaks of EBOV in the Democratic Republic of Congo underscored the unpredictable nature of ebolavirus reemergence in a region that has historically experienced outbreaks of the divergent ebolaviruses Sudan virus (SUDV) and Bundibugyo virus (BDBV)^5^. Here we show that MBP134^AF^, a pan-ebolavirus therapeutic comprising two broadly neutralizing human antibodies (bNAbs)^6,7^(see companion manuscript, Wec *et al*.) could protect against lethal EBOV, SUDV, and BDBV infection in ferrets and nonhuman primates (NHPs). MBP134^AF^ not only not only establishes a viable therapeutic countermeasure to outbreaks caused by antigenically diverse ebolaviruses but also affords unprecedented effectiveness and potency—a single 25-mg/kg dose was fully protective in NHPs. This best-in-class antibody cocktail is the culmination of an intensive collaboration spanning academia, industry and government in response to the 2013-2016 EBOV epidemic^6,7^ and provides a translational research model for the rapid development of immunotherapeutics targeting emerging infectious diseases.

---

The 2013-2016 EBOV epidemic in Western Africa and the recent EBOV outbreaks in the Democratic Republic of Congo have established ebolaviruses as pathogens of global public health relevance. Of the five ebolaviruses known to infect humans, EBOV, SUDV, and BDBV have caused outbreaks with case-fatality rates up to 90% in the last decade^5^. Although several therapeutic products are in clinical development for the treatment of Ebola virus disease (EVD), no medical countermeasures to SUDV or BDBV have progressed beyond proof-of-concept studies^1,2,4,8,9^. To address this unmet public health need, we developed a two-antibody cocktail, MBP134^AF^, with demonstrable activity against all known ebolaviruses (*companion report, Wec et al.*), including the “pre-emergent” agent Bombali virus recently discovered in molossid bats in Sierra Leone^10^. MBP134^AF^, comprising the human bNAbs ADI-15878^AF^ and ADI-23774^AF^, was selected after a systematic process including the assessment and/or optimization of multiple mAbs and their combinations for potency and breadth, Fc effector functions via glycan engineering, and *in vivo* efficacy in rodent models of EBOV and SUDV infection (*companion report, Wec et al.*). ADI-15878^AF^ and ADI-23774^AF^ both target unique, non-overlapping epitopes on the ebolavirus glycoprotein (GP), neutralize both the extracellular and endosomally cleaved forms of GP, and lack crossreactivity against the secreted GP isoform (sGP) that is abundant in the plasma of infected individuals^6,7^. The exceptional potency of MBP134^AF^ against guinea pig-adapted EBOV and SUDV (*companion report, Wec et al.*) warranted continued evaluation in the ferret and NHP large-animal models of ebolavirus challenge to assess its clinical potential.

We determined MPB134^AF^’s protective efficacy against the wild-type Makona variant of EBOV (EBOV/Makona), SUDV variant Gulu (SUDV/Gulu) and BDBV variant But-811250 (BDBV/But-811250) in the recently established ferret model, which does not require any viral adaptation and recapitulates key hallmarks of human EVD^11^-^13^. Ferrets challenged intranasally with a lethal dose of EBOV/Makona received two 15-mg doses of MBP134^AF^ three days apart, with the treatment initiated on either day 2 (ferrets 1–4) or day 3 (ferrets 5–8) post infection (p.i.) (**Fig. 1a**). MBP134^AF^ fully protected from lethal challenge in both treatment groups and cleared viremia even in ferrets 5–8, which showed signs of active infection by reverse transcription polymerase chain reaction (RT-PCR) prior to treatment (**Fig. 1b, c**). Similarly, administration of two 15-mg doses of MBP134^AF^ on days 3 and 6 (ferrets 1–4) afforded complete protection in ferrets challenged with lethal doses of SUDV/Gulu (**Fig. 1d**) or BDBV/But-811250 (**Fig. 1g**) and inhibited viral replication (**Fig. 1e-f, 1g–i**). Because SUDV/Gulu and BDBV/But-811250 have been shown to be less virulent in ferrets than EBOV/Makona, with a relatively delayed time to peak viremia^12^, we next evaluated MBP134^AF^ in a lower dose-sparing treatment course of two 5-mg doses given on days 3 and 6 p.i. In this treatment group, ferrets 5–8 challenged with SUDV/Gulu displayed high levels of viremia and uniformly succumbed to disease by day 11 p.i. (**Fig. 1d-f**). By contrast, ferrets 5–8 challenged with BDBV/But-811250 (characterized by the slowest onset of viremia^12^) were protected by the two 5-mg dose regimen, with reversion of viremia in some animals (**Fig. 1g-i).** MBP134^AF^ is the first ebolavirus therapeutic to achieve full protection in ferrets against three divergent ebolaviruses.

**Fig. 1:**
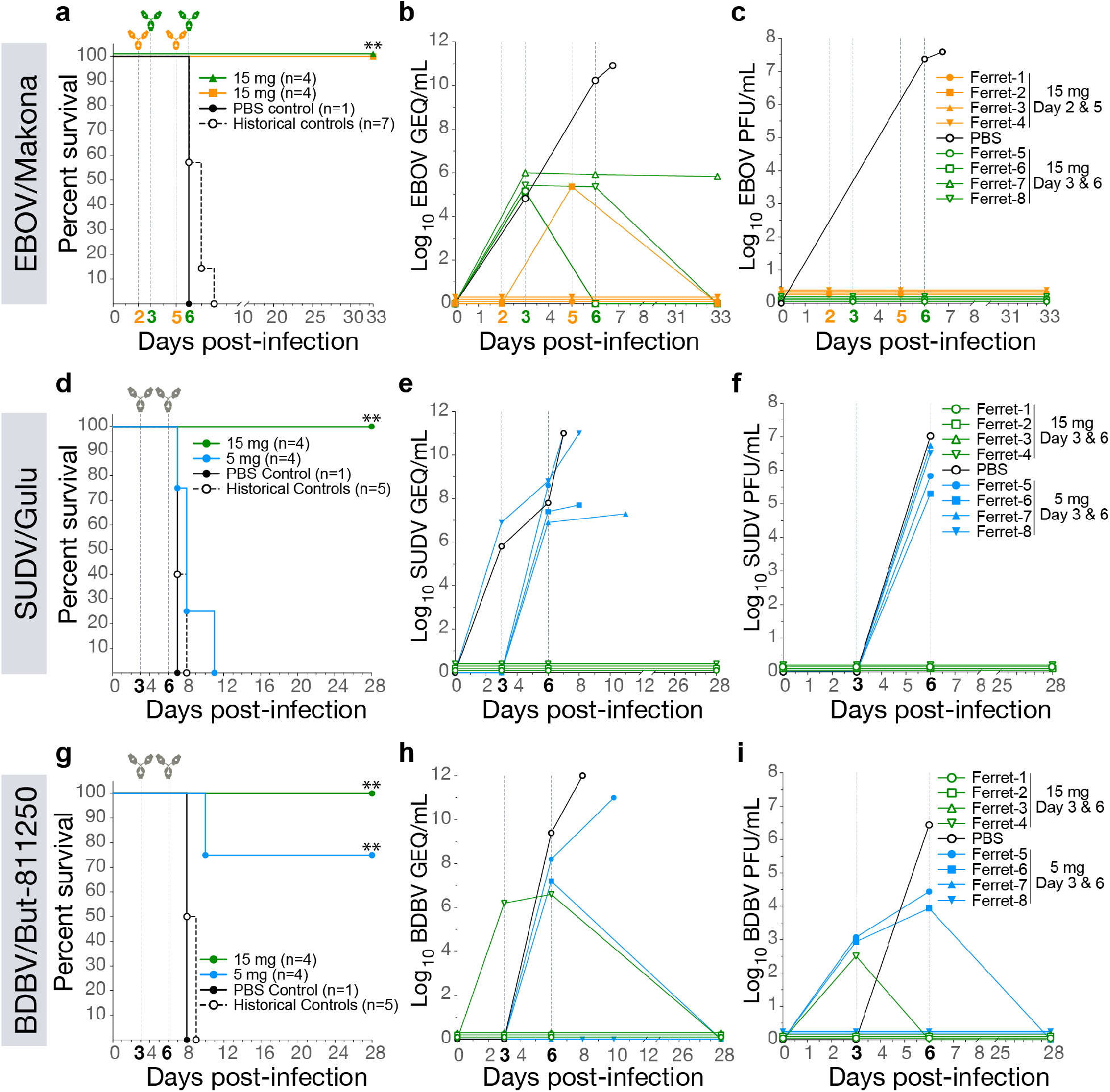
MBP134^AF^ protects ferrets from lethal EBOV, SUDV and BDBV challenge. **a,** Survival curves for ferrets challenged with EBOV/Makona and treated with 15 mg of MBP134^AF^ on either Day 2 and 5 (orange) or Day 3 and 6 (green) post-infection. **b,** Quantitative RT-PCR measuring average copies of EBOV/Makona genomic equivalents per mL of whole blood (GEQ/mL) from animals treated on day 2 and 5 (orange) and day 3 and 6 (green) post-infection. **c,** EBOV/Makona present in animals treated on day 2 and 5 (orange) or day 3 and 6 (green). **d,** Survival curves for ferrets challenged with SUDV/Gulu and treated with 15-mg (green) or 5-mg (blue) doses of MBP134^AF^ on day 3 and 6 post infection. **e,** The average SUDV/Gulu GEQ/mL from animals treated with 15- mg (green) or 5-mg doses (blue). **f,** SUDV/Gulu viremia present in animals treated with 15-mg (green) or 5-mg (blue) doses from panel D. **g,** Survival curves for ferrets challenged with BDBV/But-811250 and treated with 15-mg (green) or 5-mg (blue) doses of MBP134^AF^ on day 3 and 6 post infection. **h,** Viremia present in animals treated with two 15-mg (green) or 5-mg doses (blue) from panel G. **i,** The average BDBV/But-811250 GEQ/mL from animals treated with two 15-mg (green) or 5-mg doses (blue).

We next evaluated the MBP134^AF^ cocktail’s efficacy in the gold-standard non-human primate (NHP) model of Ebola virus challenge. Ten rhesus macaques were randomized into two treatment groups, NHPs 1–4 and NHPs 5–8, and a PBS control group of two animals, and then challenged intramuscularly (i.m.) with 1,000 plaque-forming units (PFUs) of the Kikwit variant of EBOV (EBOV/Kikwit). NHPs 1–4 received a single 25-mg/kg dose of MBP134^AF^ on day 4 p.i., whereas NHPs 5–8 received a more conservative two-dose regimen of 50 mg/kg then 25 mg/kg on days 4 and 7 p.i., respectively. Remarkably, the single 25-mg/kg dose of MBP134^AF^ completely reversed the onset of EVD and protected NHPs 1–4 from a lethal EBOV/Kikwit exposure (**Fig. 2a**). All animals in this study were confirmed to have had an active EBOV/Kikwit infection via RT-PCR (10^7^–10^10^ viral genome equivalents per mL (GEQ/mL)) and plaque assay (10^3^–10^6^ PFU/mL) prior to treatment on day 4 p.i. (**Fig. 2b, c**). These high levels of viremia could nonetheless be reversed by MBP134^AF^ treatment—viremia in animals from both treatment groups fell below the limit of detection in the plaque assay by day 7 p.i. and in the RT-PCR assay by day 14 p.i. (**Fig. 2b, c**). Fever was detected in control animals and in three out of four animals in each treatment group at the time of the first MBP134^AF^ dosing; however all treated animals returned to normal body temperature by day 10 p.i. Treated animals also maintained substantially lower clinical scores and reduced grade of thrombocytopenia compared to control NHPs (**Fig. 2d-f**). Two animals (NHP-3 and NHP- 8), one from each treatment group, showed significant signs of EVD-induced liver injury prior to treatment, with elevated C-reactive protein (CRP), alanine aminotransferase (ALT) and aspartate aminotransferase (AST). These and other hallmarks of EVD were significantly reduced post-treatment with MBP134^AF^ by day 10 p.i. (**Fig. 2d-i, Extended Data Fig. 1**). Thus, the pan-ebolavirus MBP134^AF^ cocktail could potently reverse the course of EVD and deliver complete therapeutic protection in NHPs following a lethal EBOV/Kikwit challenge with a single dose of only 25 mg/kg.

**Fig. 2:**
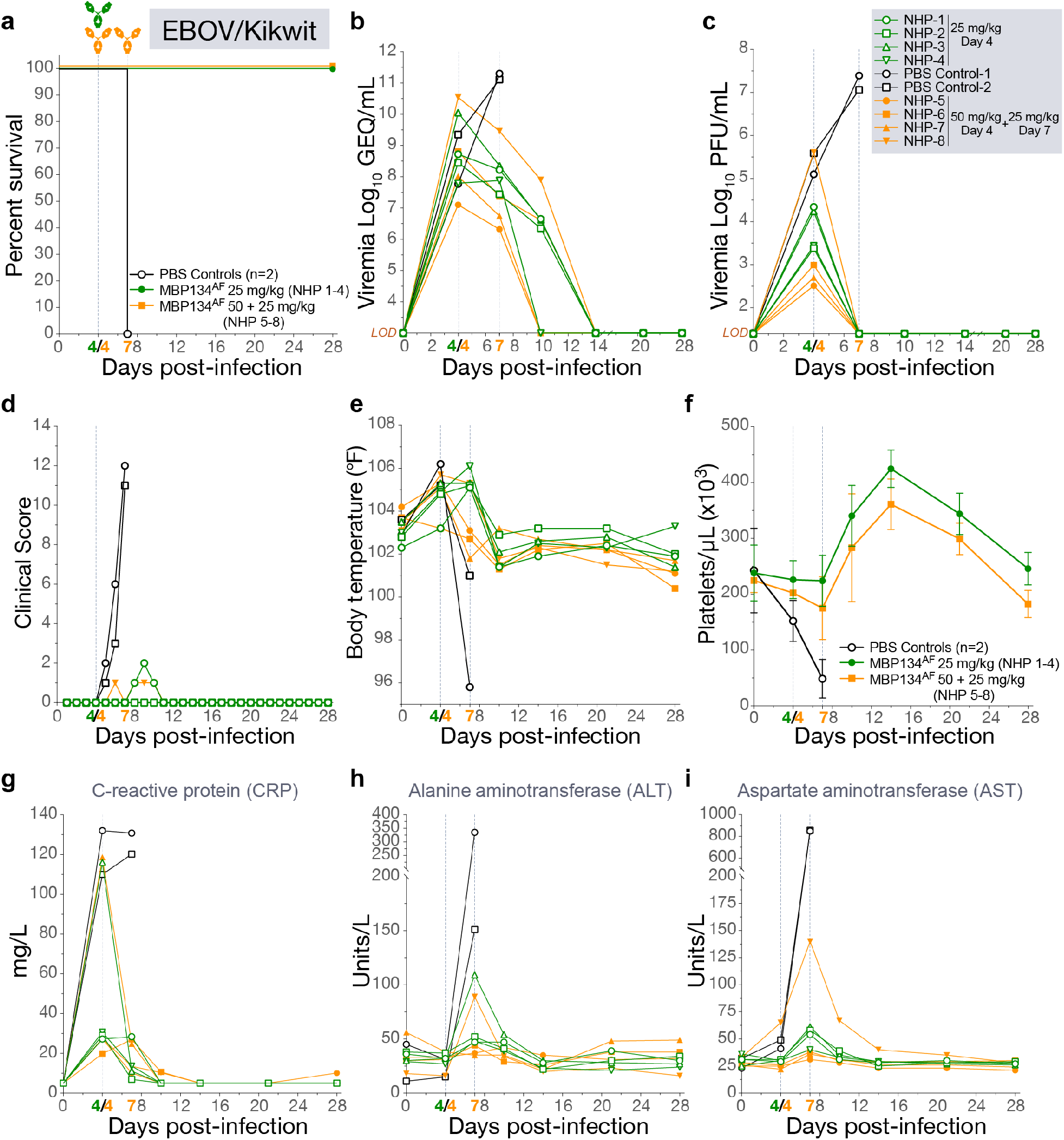
A single 25 mg/kg dose of MBP134^AF^ protects rhesus macaques challenged with EBOV/Kikwit. **a,** Survival curves for NHPs challenged with EBOV/Kikwit and treated with a single 25- mg/kg dose of MBP134^AF^ on day 4 (green) post-infection or a more conservative two-dose regimen of 50 mg/kg on day 4 and 25 mg/kg on day 7 (orange) post infection. **b,** The average EBOV/Kikwit GEQ/mL from animals treated with a single dose of MBP134^AF^(green) or two doses of MBP134^AF^(orange). All detectable EBOV/Kikwit was eliminated 10 days post treatment on day 4 post infection. **c,** Plaque-forming units (PFU) of infectious EBOV/Kikwit present in animals treated with either a single (green) or two-dose course of MBP134^AF^(orange). Infectious EBOV/Kikwit was no longer detectable by plaque assay by the next bleed of treated animals on day 7 post infection. **d,** Clinical scores of animals within the study cohort are shown. Aside from the control animals only NHP-1 and NHP-8 scored for partially or completely refusing their daily nutrition. **e,** Body temperatures taken during the course of the study show 9 of the ten animals registered an elevated temperature by day 7 p.i. Animals receiving MBP134^AF^ returned to baseline temperature by day 10 p.i. **f,** The platelet counts for the cohort show severe thrombocytopenia in the control animals post-infection. By contrast, animals receiving MBP134^AF^ rapidly recovered and displayed little or no signs of thrombocytopenia. **g,** The graphed C-reactive protein (CRP) levels show NHP-3 and NHP-8 suffered from acute inflammation as a result of EVD prior to MBP134^AF^ treatment. **h,** The alanine aminotransferase (ALT) levels from each animal are shown, NHP-3 and NHP-8 again demonstrate signs of advanced EVD that were alleviated post-treatment. **i,** Aspartate aminotransferase (AST) levels for all the challenged animals shows initial signs of liver damage resultant of EVD. Samples from NHP-8 in particular show a significant spike in AST levels that returned to baseline post-treatment. Legend for graphs in top right-hand corner, *LOD* = limit of detection.

In the experiments described above, MBP134^AF^ was produced using a *Nicotiana benthamiana* (tobacco) plant-based expression system^14,15^. However, because the manufacturing infrastructure for Nicotiana-based products is still limited, we sought to transition MBP134^AF^ to the well-established Chinese hamster ovary (CHO) cell production platform. Accordingly, we expressed MBP134^AF^ in a GDP-fucose transporter SLC35C1-knockout cell line (CHOK1-AF), which maintains the afucosylated state of MBP134^AF,16^. Comparative studies indicated that CHOK1-AF–produced MBP134^AF^ is comparable or even surpasses its plant-produced counterpart in neutralization potential (data not shown), Fc effector functions relevant to this cocktail’s antiviral potency (**Extended Data Fig. 2**), and protective efficacy in guinea pigs (**Extended Data Fig. 3**). Therefore, the Nicotiana- and CHO-produced MBP134^AF^ products are functionally equivalent. Accordingly, all remaining experiments described herein were performed with CHOK1-AF expressed MBP134^AF^, the manufacturing system being employed for its clinical development.

We tested MBP134^AF^ in a blinded NHP study in which rhesus macaques were challenged i.m. with SUDV variant Boniface (SUDV/Boniface; 1,000 PFU). This model typically affords 50% lethality^17^(unpublished data). Twelve animals were randomized into two treatment groups, NHPs 1–4 and NHPs 5–8, and one control group, PBS controls 1–4. On day 5 p.i., NHPs 1–4 and 5–8 received single 7.5-mg/kg and 25-mg/kg doses of MBP134^AF^, respectively. Both doses of MBP134^AF^ provided full protection from SUDV disease and all MBP134^AF^-treated animals became viremia-negative by day 8 p.i. (**Fig. 3a-c**). MBP134^AF^-treated animals displayed little to no clinical signs of disease in contrast to the control animals—half of the latter succumbed to infection (**Fig. 3d**). Importantly, all of the animals in the blinded MBP134^AF^ cohort registered a fever prior to receiving MBP134^AF^ on day 5 p.i. (**Fig. 3e**). The two surviving control animals remained viremic past day 14 p.i. (**Fig. 3b, c**), maintained elevated ALT, AST, and alkaline phosphatase (ALP) levels, and showed significant thrombocytopenia (**Fig. 3f-i, Extended Data Fig. 4**) out to day 21 p.i.

**Fig. 3:**
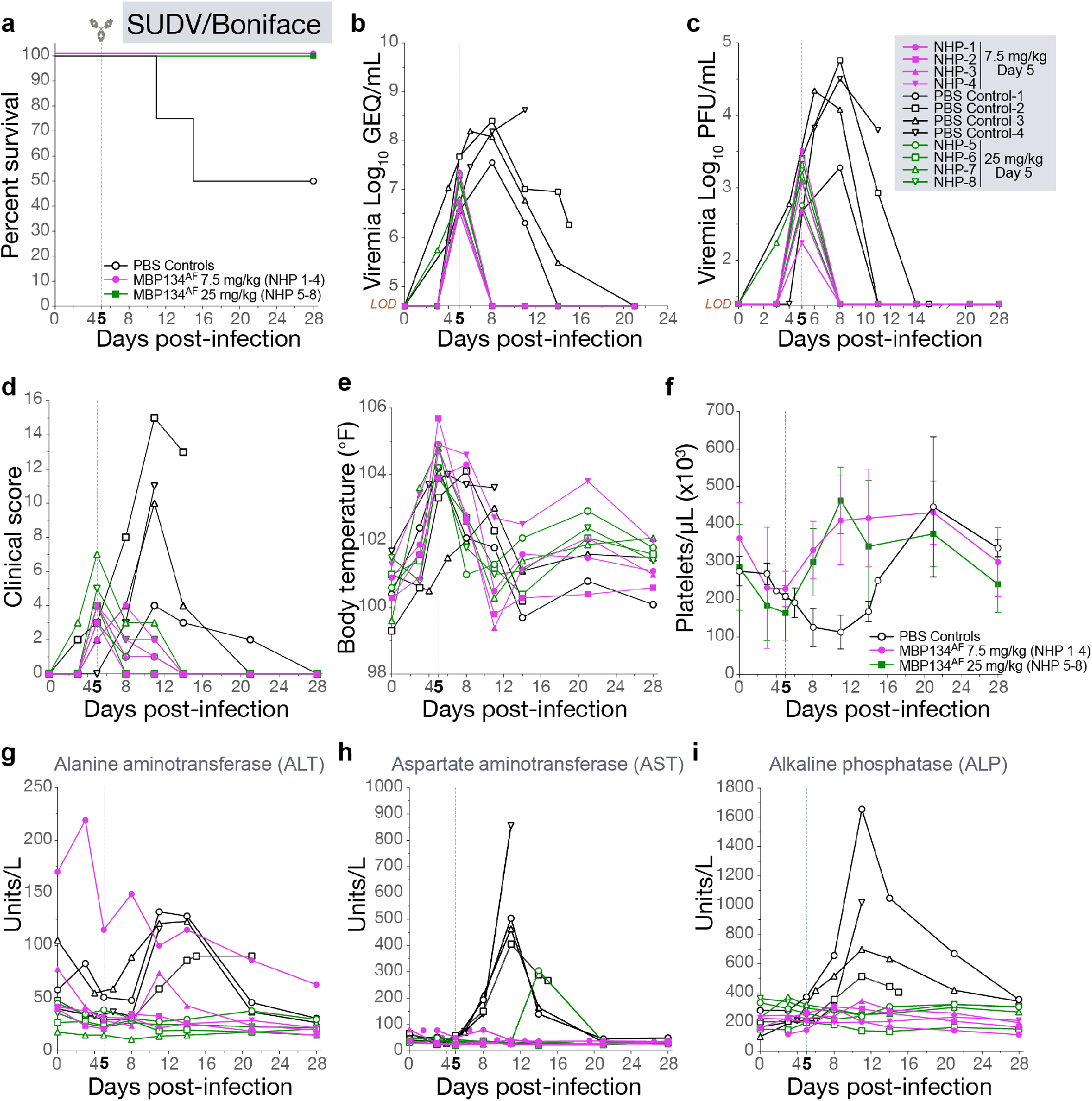
A single 25 mg/kg or 7.5 mg/kg dose of MBP134^AF^ protects rhesus macaques challenged with SUDV/Boniface. **a,** Survival curves for NHPs challenged with SUDV/Boniface receiving either PBS (black), a 25-mg/kg dose of MBP134^AF^(green) or a 7.5-mg/kg (purple) dose of MBP134^AF^ on day 5 post infection. **b,** The average SUDV/Boniface GEQ/mL from animals receiving a 25- mg/kg dose of MBP134^AF^(green) or a 7.5-mg/kg dose of MBP134^AF^ (purple). All detectable SUDV/Boniface was eliminated by day 8 post infection (the next bleed post treatment). **c,** Plaque-forming units (PFU) of infectious SUDV/Boniface present in animals receiving a 25-mg/kg dose of MBP134^AF^(green) or a 7.5-mg/kg dose of MBP134^AF^(purple) as well as in PBS controls (black). Infectious SUDV/Boniface was no longer detectable by plaque assay by the next bleed on day 8 p.i. **d,** All of the animals in the cohort had clinical scores on day 5 p.i. prior to receiving MBP134^AF^. Animals dosed with MBP134^AF^ no longer registered a clinical score by day 14 p.i., while the surviving controls continued scoring out to day 21 p.i. **e,** Body temperatures taken from the blinded cohort showed all the animals registered an elevated temperature by day 5 p.i., prior to receiving MBP134AF. **f,** The platelet counts for the cohort show declining counts p.i. with the MBP134^AF^ treated animals rapidly recovering and the PBS controls displaying severe thrombocytopenia. **g,** The alanine aminotransferase (ALT) levels from each animal are shown; the control animals all display elevated levels from baseline post infection. In contrast, all of the animals receiving MBP134^AF^ show little to no signs of EVD. Graph legend in panel F. **h,** The aspartate aminotransferase (AST) levels for all the MBP134^AF^ treated animals show little to no increase in AST levels, while the control animals all show highly elevated AST levels by day 10 post-infection. **i,** The alkaline phosphatase (ALP) levels are graphed for each animal in the cohort, no sign of EVD induced ALP levels are present in any of the MBP134^AF^ treated animals. In contrast, all of the control animals that received PBS display elevated ALP levels resulting from EVD. Legend for graphs in top right-hand corner, *LOD* = limit of detection.

We next determined the protective efficacy of MBP134^AF^ against lethal BDBV/But-811250 challenge in the cynomolgus macaque model of infection^5,9^. We chose to treat animals at day 7 p.i. with MBP134^AF^ because previous reports indicated that they were already viremic and showing signs of EVD at this time point^9^. We reasoned that treatment under these post-exposure conditions would afford a rigorous evaluation of MBP134^AF^’s ability to reverse advanced EVD caused by BDBV.

Accordingly, a cohort of 9 animals was exposed to 1,000 PFU (i.m.) of BDBV/But-811250. Six randomly selected animals received a single 25-mg/kg i.v. infusion of MBP134^AF^ on day 7 p.i. and three received PBS. This single dose of MBP134^AF^ provided significant levels of protection (P value of 0.006 or 0.0108 if calculated including the historical controls^9^), with only one animal succumbing to infection. By contrast, uniform lethality was observed in the PBS control group (**Fig. 4a**). Prior to MBP134^AF^ treatment on day 7 p.i., animals registered elevated clinical scores and body temperatures, viremia as high as ∼10^11^ GEQ/mL or 10^7^ PFU/mL, and EVD-induced thrombocytopenia (**Fig. 4b-f**). By the next blood collection point, on day 10 p.i., animals that received MBP134^AF^ had no detectable infectious BDBV in the blood, and clinical scores were reduced to basal levels by day 12 p.i., a complete reversion of infection and disease. The single treated animal that succumbed, NHP-5, did not have the highest viral load but showed acute liver injury prior to treatment, displaying the highest ALP, ALT, and AST levels of all the animals in the cohort on day 7 p.i. prior to receiving its dose of MBP134^AF^(**Fig. 4g-i, Extended Data Fig. 5**). Given the recovery of two animals (NHP-4 and NHP-6) harboring higher viral loads prior to treatment, we postulate that NHP-5’s liver injury prior to treatment was too severe for it to recover despite receiving MBP134^AF^. To our knowledge, MBP134^AF^ is the first therapeutic to demonstrate significant levels of protection and reversion of BDBV disease in cynomolgus macaques.

**Fig. 4:**
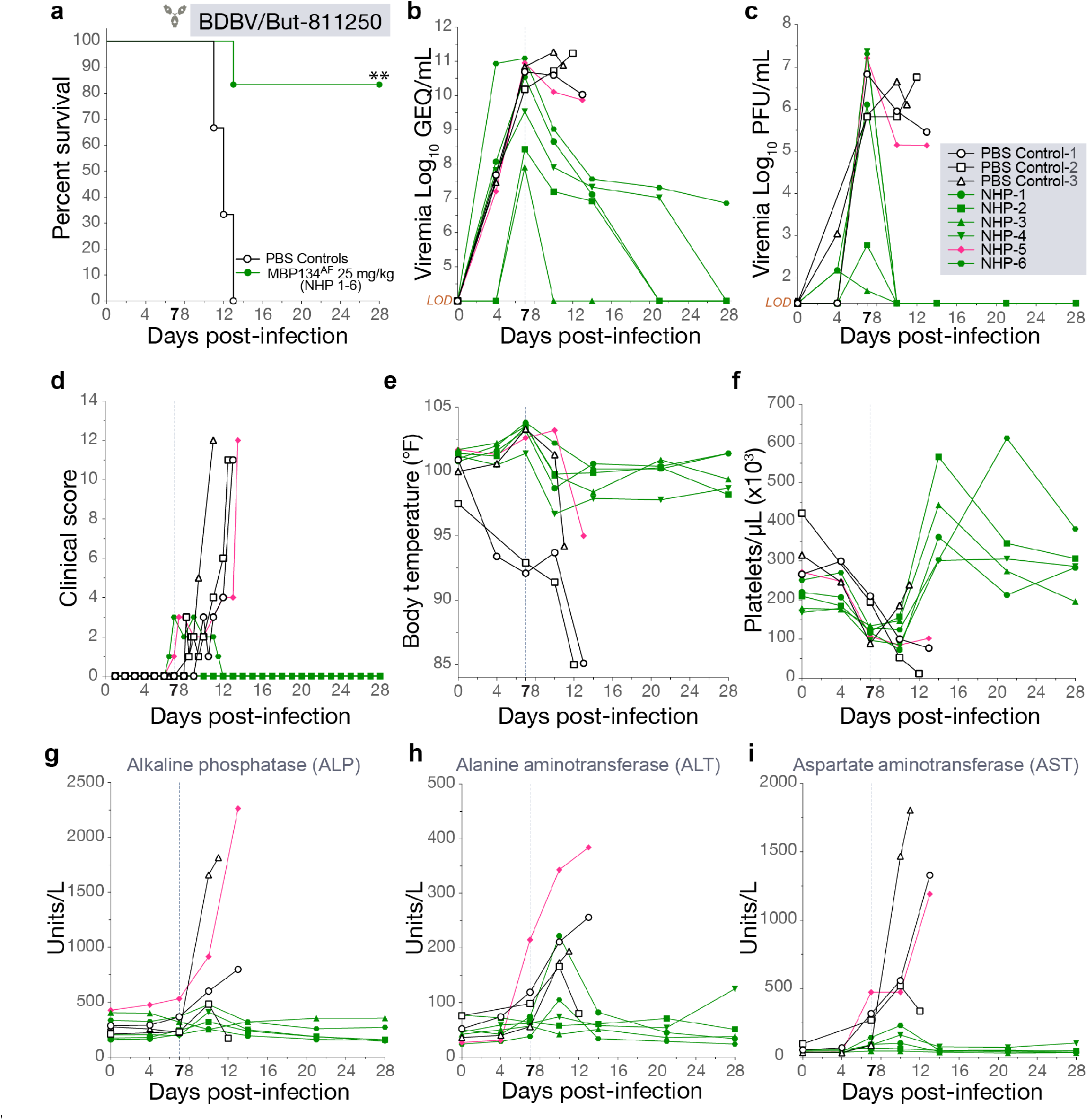
A single 25 mg/kg dose of MBP134^AF^ protects cynomolgus macaques challenged with BDBV/But-811250. **a,** Survival curves for NHPs challenged with BDBV/But-811250 and treated with a single 25-mg/kg dose of MBP134^AF^ on day 7 post infection (green) or PBS (black). **b,** The average BDBV/But-811250 GEQ/mL from animals treated with a single dose of MBP134^AF^(green) or PBS (black). Serum samples taken from NHP-4 and NHP-6 on days 20 and 28 tested negative for viral genetic material. **c,** Plaque-forming units (PFU) of infectious BDBV/But-811250 present in animals treated with MBP134^AF^(green) or PBS (black). Infectious BDBV was no longer detectable by plaque assay by the next bleed on day 10 post infection in the surviving animals. **d,** Clinical scores show NHP-5 and NHP-6 registering scores from refusing nutrition, having a hunched posture and displaying petechiation over 10% of their bodies prior to receiving MBP134^AF^ on day 7 p.i. While NHP-6 cleared clinical signs of infection by day 12 p.i., NHP-5 failed to recover and succumbed to infection on 13 p.i. **e,** Body temperatures from all the animals showed the majority registered an elevated temperature by day 7 p.i. prior to receiving MBP134^AF^. **f,** The platelet counts for the cohort show thrombocytopenia occurring in all the animals by day 7 p.i. prior to receiving MBP134^AF^. All the MBP134^AF^ treated animals (excluding NHP-5) cleared signs of thrombocytopenia by day 14 p.i., seven days post treatment. **g,** The alkaline phosphatase (ALP) levels are graphed for each animal in the cohort. **h,** The alanine aminotransferase (ALT) levels from each animal are shown. Notably NHP-5 displayed advanced signs of EVD prior to treatment. **i,** Aspartate aminotransferase (AST) levels for all the challenged animals shows initial signs of liver damage resultant of EVD. Samples from NHP-5 in particular show a significant spike in AST levels suggesting a severe onset of disease beyond that of the other animals in the cohort, a possible explanation of NHP-5’s succumbing to EVD despite receiving the MBP134^AF^ treatment course. Legend for graphs in top right-hand corner, *LOD* = limit of detection.

Prior to this work, the development of monoclonal antibody-based therapeutics has typically followed a “one bug, one drug” paradigm under the premise that mAbs with broad activity would not be as potent as those with clade-specific activity^18^. Here, we demonstrate that MBP134^AF^, a pan-ebolavirus immunotherapeutic comprising two bNAbs ADI-15878^AF^ and ADI-23774^AF^ could not only protect NHPs against every ebolavirus known to cause human disease outbreaks but could do so at an unparalleled single 25- mg/kg dose. Importantly, MBP134^AF^ was effective against multiple ebolaviruses from different species, suggesting that it will retain activity in the face of both intra- and inter-species sequence divergence, a result of its targeting highly conserved epitopes in GP. Indeed, as shown in the companion paper *(Wec et al.)*, MBP134^AF^ recognizes and neutralizes entry by the newly identified Bombali virus glycoprotein. Further studies exploring 7.5 mg/kg or lower intravenous doses of MBP134^AF^ could open the door to intramuscular or subcutaneous delivery via autoinjector, allowing for rapid and efficient drug administration to patients and reducing the burden on healthcare workers in the field and Ebola virus treatment units. The developmental path of MBP134^AF^ presents a model for the rapid design of next-generation antiviral immunotherapeutics targeting World Health Organization-priority pathogens.

## Acknowledgements

This project has been funded in part with federal funds from the Department of Defense under Contract No. HDTRA1-13-C-0018; the Department of Health and Human Services’ National Institutes of Health (U19AI109711, U19AI109762, AI1332204 and AI132256), the Public Health Agency of Canada (PHAC), and Office of the Assistant Secretary for Preparedness and Response, and Biomedical Advanced Research and Development Authority (BARDA), under Contract No. HHSO100201700023C. K.C. was additionally supported by an Irma T. Hirschl/Monique Weill-Caulier Research Award. Opinions, conclusions, interpretations, and recommendations are those of the authors and are not necessarily endorsed by the US. Army. The mention of trade names or commercial products does not constitute endorsement or recommendation for use by the Department of the Army or the Department of Defense.

## Contributions

L.Z., K.C., Z.A.B., X.Q., T.W.G., and J.M.D. conceived the overall study. A.S.H., C.E.M, S.H., R.W.C, J.B.G., V.B., R.M.J., M.N.R, W.Z, L.B, K.T, X.Q., T.W.G and J.M.D. performed the *in vivo* studies and data analyses in Figs. 1-4 and Extended Data Figs. 1, 3-5. D.M.A and Z.A.B. prepared MBP134^AF^ for the ferret and guinea pig studies. O.B., N.B., J.V., M.P., and K.J.W., all contributed to the development and expression of the plant derived MBP134^AF^. W.S.S. and E.A. developed the CHOK1-AF clonal pools for MBP134^AF^. D.K. manufactured and formulated the CHOK1-AF expressed MBP134^AF^ for the NHP studies. A.Z.W. carried out VSV-based neutralization experiments to verify activity CHOK1-AF mAb lots prior to the NHP studies. B.G. and G.A. carried out mAb effector function studies reflected in Extended Data Fig. 2. L.Z., K.C., A.Z.W., Z.A.B., A.S.H., C.E.M., S.H., X.Q., T.W.G., and J.M.D. wrote the manuscript with contributions from all of the authors.

